# Alfalfa leaf curl virus is transmitted by *Aphis craccivora* in a highly specific circulative manner

**DOI:** 10.1101/832204

**Authors:** F. Ryckebusch, N. Sauvion, M. Granier, P. Roumagnac, M. Peterschmitt

## Abstract

Two members of the genus Capulavirus (Geminiviridae) are transmitted by aphids including Alfalfa leaf curl virus (ALCV) transmitted by *Aphis craccivora*. The capulavirus Euphorbia caput-medusae latent virus was shown here to be transmitted also by *A. craccivora*, using the population EuphorbiaSA. ALCV was transmissible by several *A. craccivora* populations including Robinia, but not EuphorbiaSA, reflecting a high transmission specificity. ALCV accumulation and localization were analyzed in whole insects, midguts, hemolymphs, and heads of aphids from both populations and from the non-vector species *Acyrthosiphon pisum*. A 6-day persistence was observed in *A. craccivora* populations but not in *A. pisum*. Vector and non-vector *A. craccivora* populations could be distinguished by contrasted virus accumulations and midgut intracellular localization. Results confirm that ALCV is transmitted according to a circulative-persistent mode, and are consistent with a gut barrier to the transmission of ALCV in *A. pisum* and a primary salivary gland barrier in *A. craccivora*.

## 1. Introduction

Nearly 55% of reported plant pathogenic viruses are transmitted by piercing-sucking insects of the Hemiptera order (Hogenhout et al., 2008). In the viral family *Geminiviridae*, a large range of hemipterans were identified as vectors. Geminiviruses of the genus Begomovirus are transmitted by whiteflies (i.e., Aleyrodidae) (Cohen and Nitzany, 1966), viruses of the genera Becurtovirus, Curtovirus, Mastrevirus, Turncurtovirus are transmitted by leafhoppers (i.e, Cicadellidae) (Heydarnejad et al., 2013; Horn et al., 1994; Briddon et al., 1992; Razavinejad et al., 2013) and those belonging to the genera Grablovirus and Topocuvirus are transmitted by treehoppers (i.e., Membracidae) (Bahder et al., 2016; Briddon et al., 1996). Interestingly, although Aphididae is the insect group with the highest number of species reported as vector of plant viruses, aphid transmission was discovered only in 2015 when aphids of the species *Aphis craccivora* Koch, 1854 (Aphididae) were shown to transmit Alfalfa leaf curl virus (ALCV), a geminivirus belonging to the newly defined genus Capulavirus (Roumagnac et al., 2015; Varsani et al., 2017). Since then, Plantago latent virus (PlLV) another capulavirus was shown to be transmitted by the aphid *Dysaphis plantaginea* (Passerini, 1860) (Susi et al., 2019) suggesting that aphid transmission is a taxonomic criterion of the new genus which comprises two additional members, Euphorbia caput-medusae Latent Virus (EcmLV) and French bean Severe Leaf Curl Virus (FbSLCV).

Contrasted transmission specificities were observed between genera within the family *Geminiviridae*. While begomoviruses, irrespective of their geographic origin, are transmitted by whiteflies of the species complex *Bemisia tabaci* (Bedford et al., 1994; De Barro et al., 2011), mastreviruses are transmitted by leafhoppers of various genera depending on their geographic origin. Therefore, considering the scarcity of transmission data about capulaviruses, their transmission specificity is quite unpredictable. While it is clear that unlike the begomovirus genus, there are more than one vector species associated with the capulavirus genus, it was not known how far these capulavirus vectors are specific in their ability to transmit capulaviruses.

The first objective of this study was to further support that aphids are vectors of capulaviruses and thus confirm that it may be a taxonomic criterion of the new genus. The second objective was to assess the specificity of the aphid transmission. The identification of an aphid vector of a third capulavirus (EcmLV) confirmed that aphid transmission can be considered as a taxonomic criterion of this new genus. Moreover, the transmission tests of ALCV performed with various aphid species and populations showed that only aphids of the *A. craccivora* group transmitted ALCV, and most importantly, not all *A. craccivora* populations were detected to be vector. According to viral DNA persistence monitored with qPCR in vector and non-vector aphids, and its localization by FISH, the barrier associated with the very high transmission specificity is located at the salivary gland level. A second transmission barrier was detected at the gut level.

## 2. Material and Methods

### 2.1. Virus inoculation and detection in plants

#### 2.1.1. Preparation of agroinfectious clones and agroinoculation

The agroinfectious clones of EcmLV and ALCV were reported previously (Bernardo et al., 2013, Roumagnac et al., 2015). An agroinfectious clone of the reported PILV clone (Genbank accession number KT214390) (Susi et al., 2017) was prepared as follows. Its genome was released from its Pjet1.2 vector by *Pst*I restriction and ligated as a tandem repeat into the corresponding restriction site of the binary vector pCambia 2300. *Agrobacterium tumefaciens* strain C58-MP90 was transformed with the recombinant plasmid by electroporation. Agrobacteria were grown overnight at 28°C in LB medium containing gentamicin and kanamycin until an optical density (OD600) of 2-3. Bacterial suspensions were then centrifuged at 1000 g for 25 min. The pellets were resuspended in ultrapure water (Milli-Q) containing MgCl2 (10mM) and acetosyringone (150mM). Agroinoculations were performed with a syringe by repeated needle injections at the base of the stem on ten-day-old broad bean plants (*Vicia faba*, cv. ‘Sevilla’), 14-day old tomato plants (*Solanum lycopersicum* L.), one-month-old buckhorn plantain (*Plantago lanceolata* L.) or 6-year old euphorbia medusa’s head plants (*Euphorbia caput-medusae* L.).

#### 2.1.2. DNA extraction

The infection of ALCV, EcmLV, and PlLV inoculated plants was monitored 4 to 6 weeks after inoculation by symptom observation (ALCV) and/or by PCR-mediated detection of viral DNA (ALCV, EcmLV, and PlLV) in total plant DNA extracts. From each plant, four leaf disks of 4 mm diameter were collected from the four youngest leaves, one disc per leaf, and stored at – 20°C. Total DNA was extracted by grinding leaf disks in 400 µL of modified Edwards buffer containing 200 mM Tris-HCl (pH 7.5), 25 mM EDTA, 250 mM NaCl, 0.5% SDS, 1% PVP40 and 0.2% ascorbic acid. The extract was incubated at 65°C for 10 min and centrifuged at 15700 g for 10 min. One volume of isopropanol was added to the supernatant before a 20 min centrifugation at 15700 g. After resuspension of the pellet in 500 µL of 70% ethanol, nucleic acids were recovered by centrifugation (15700 g, 15 min) and resuspended in 50 µL sterile distilled water. The DNA extracts were stored at −20°C before use.

#### 2.1.3. PCR detection of viral DNA

PCR primers designed to detect ALCV and EcmLV prime the amplification of a 174 bp fragment from the Rep gene: ALCV2cEcmLV-F, 5’-GAG GAA TTC GGA CTT GGA TG -3’ and ALCV2cEcmLV-R, 5’-TTC TTC GAC ATC AAG GAC CC -3’. PCR primers designed to detect PlLV prime the amplification of a 293 bp fragment from the CP gene: PlLV_729-F, 5’-AAG GGA AAG GCT GGT TAT GG -3’ and PlLV_1013-R, 5’-GAA TCT CTT CTC TGA ATC GTG GTC -3’. The cycling protocol for all primers was as follow: 2 min denaturation at 95°C, 1 min primer annealing at 60°C and 50 sec DNA extension at 72°C followed by 30 cycles each consisting of 1 min at 94°C, 1 min at 60°C, and 50 sec at 72°C. The PCR program was terminated by a 5 min incubation at 72°C. The PCR products were resolved by electrophoresis in a 1% agarose gel and ethidium bromide staining.

### 2.2. Species and populations of aphids

#### 2.2.1. Origin of aphids

The aphids used for transmission tests were from rearings initiated with aphids collected in France, South Africa, or Switzerland. Aphids of the following species were from France: *A. craccivora*; *D. plantaginea*; *Acyrthosiphon pisum* (Harris, 1776); *Aphis fabae* Scopoli, 1763; *Aphis gossypii* Glover, 1877; *Myzus persicae* (Sulzer, 1776); *Therioaphis trifolii* (Monell, 1882). Full description of aphids is provided in Supplemental Table S1. *A. craccivora* aphids were from three populations collected on Fabaceae species near Montpellier (France), namely *Robinia pseudoacacia* L. (false acacia), *Vicia sativa* (L.) Bernh. (common vetch), and *Medicago sativa* L. (alfalfa). These populations were respectively named population Robinia, Vicia, and Medicago. The favorite vector candidate for a potential aphid transmission of EcmLV was a black-backed aphid observed on euphorbia Medusa’s head plants, the natural host of EcmLV. A rearing of this aphid was established in a P3 containment chamber with individuals collected in 2015 in the Buffelsfontein Game and Nature Reserve (Darling region of the Western Cape, South Africa) where EcmLV was detected for the first time (Bernardo et al., 2013). It was identified here with molecular and morphological criteria. Taxonomy and nomenclature were as described by Remaudière and Remaudière, 1997, Blackman and Eastop, 2000, and Favret, 2014.

#### 2.2.2. Molecular identification of aphids

The total DNA was purified from individual aphids using the CTAB method as described in Peccoud et al., 2013. DNA was recovered in 50 µl of ultra-pure H_2_O. A 658-bp fragment of the mitochondrial cytochrome c oxidase subunit I gene (COI) was amplified with a mix of degenerate primers adapted from the LCO1490/HCO2198 universal primers (Folmer et al., 1994) to Sternorrhyncha (Isabelle Meusnier, com. pers.), the suborder of the Hemiptera that include Aphididae. They consist of two forward primers namely LCO1490stern1_t1 (5’-TGTAAAACGACGGCCAGTTTASAACTAACCACAAARMTATTGG-3’), LCO1490stern2_t1 (5’-TGTAAAACGACGGCCAGTTTTCAACTAATCATAARGATATTGG-3’), and two reverse primers namely HCO2198Stern1_t1 (5’-CAGGAAACAGCTATGACTAWACTTCWGGATGTCCAAAAAAYCA-3’), and HCO2198Stern2_t1 (5’-CAGGAAACAGCTATGACTAMACCTCAGGATGHCCAAAAAATCA-3’). The PCR was performed in a final volume of 30 µL containing: 3 µL of QIAGEN Coraload buffer (containing 45 pmol MgCl2), 0.1 mM of each dNTP, 0.5 mM of MgCl2 (Qiagen), 0.2 µM of each primer, 0.625 U of Taq DNA Polymerase (Qiagen) and 2 µL of DNA extract. The PCR cycles were as follows: initial denaturation at 94°C for 2 min; followed by 5 cycles at 94°C for 30 s, 45°C for 40 s and 72°C for 1 min, and then 35 cycles at 94°C for 30 s, 51°C for 40 s and 72°C for 1 min, with a final 10-min extension period at 72°C. PCR products were purified and Sanger-sequenced in both directions by Beckman Coulter Genomics (Takeley, UK) with M13 primers complementary to the 5’ ends of the PCR primers. Chromatograms were aligned by the Muscle algorithm, cleaned and visually checked under Geneious Pro 10.2.6 (http://www.geneious.com). All sequences were deposited in GenBank. Sequences were aligned with Genbank or BOLD (http://www.barcodinglife.org) COI sequences from aphids of the *A. craccivora* group - from different geographical origins-, and of other species, in particular species previously described as closely related to the *A. craccivora* group (e.g. *Aphis coronillae* Ferrari, 1872; *Aphis intybi* Koch, 1855; *Aphis lhasaensis* Zhang, 1981). Aphids from which COI sequences were downloaded from Genbank or BOLD are described in Supplemental Table S2. A neighbor joining tree (Saitou and Nei, 1987) was constructed from Jukes and Cantor’s distance matrix using the PAUP* 4.0b10 (Swofford, 2001) plugin in Geneious. Node support was calculated from 10,000 bootstrap replicates.

### 2.3. Transmission tests

Broad bean, buckhorn plantain, and alfalfa plants were kept in P2 containment chambers under 16 h light at 26±2 °C, and 8 h dark at 24±2 °C. Tomato and euphorbia medusa’s head plants were maintained in a P3 containment chamber with the same temperatures but with 14 h light. The duration of the acquisition access period (AAP) was adapted to each test (Table 1) and carried out with 50 aphids per source plants. The duration of the inoculation access period (IAP) and the number of insects transferred to each test plant for the IAP were also adapted to each test. The IAP was stopped by spraying the test plants with the insecticide Pirimor G (1g/L in water).

**Table 1.** Transmission tests of three capulaviruses with aphids of various species. The rate of infected plants is in bold for successful transmissions.

| Virus species | Aphid species | Rate of infected plants | Host plant | AAP (days) | IAP (days) | Number aphids / IAP plant |
| --- | --- | --- | --- | --- | --- | --- |
| PILV | *Dysaphis plantaginea* Nov08 | **9/10** | *Plantago lanceolata* | 27 | 5 | 10 |
|  | *Aphis gossypii* NM1 | 0/20 | *Plantago lanceolata* | 13 | 5 | 10 |
|  |  | 0/20 |  | 21 | 5 | 10 |
| EcMLV | *Aphis craccivora* EuphorbiaSA | **2/2** | *Euphorbia caput-medusae* | weeks | 35 | 50 |
|  | *Aphis gossypii* NM1 | 0/9 | *Solanum lycopersicum* | 2 | 5 | 10 |
|  | *Myzus persicae* Lav85 | 0/24 | *Solanum lycopersicum* | 2 | 5 | 10 |
|  | *Dysaphis plantaginea* Nov08 | 0/12 | *Plantago lanceolata* | 3 | 3 | 10 |
| ALCV | *Aphis craccivora* Robinia | **8/15** | *Vicia faba* | 2 | 5 | 10 |
|  |  | **9/30** |  | 2 | 5 | 10 |
|  | *Aphis fabae* A06-405 | 0/19 | *Vicia faba* | 2 | 5 | 5 |
|  | *Aphis fabae* AFCOL | 0/3 | *Vicia faba* | 2 | 5 | 5 |
|  |  | 0/5 |  | 2 | 5 | 5 |
|  |  | 0/10 |  | 2 | 5 | 10 |
|  | *Aphis gossypii* NM1 | 0/20 | *Vicia faba* | 2 | 5 | 5 |
|  | *Myzus persicae* Lav85 | 0/6 | *Medicago sativa* | 2 | 5 | 20 |
|  | *Acyrtosiphon pisum* LL01 | 0/20 | *Vicia faba* | 2 | 5 | 10 |
|  | *Therioaphis trifolii* Mon18 | 0/1 | *Medicago sativa* | 15 | 5 | 10 |
|  | *Aphis craccivora* EuphorbiaSA | 0/45 | *Vicia faba* | 2 | 5 | 10 |
|  |  | 0/15 |  | 2 | 5 | 10 |
|  |  | 0/6 |  | 3 | 5 | 30-150 |
|  |  | 0/12 |  | 7 | weeks | 10 |

#### 2.3.1. ALCV transmission

Aphids of *A. fabae, A. gossypii, M. persicae, A. pisum, T. trifolii* and four populations of *A. craccivora* (Robinia, Medicago, Vicia, and EuphorbiaSA), were tested for their ability to transmit ALCV. The virus acquisition feeding was performed on broad bean plants 4 to 6 weeks after their agroinfection with ALCV, except one test in which it was performed on naturally infected alfalfa. Virus inoculation was performed on 8-day-old broad bean or one-month-old alfalfa plants. The transmission success was assessed by symptom observation and detection of ALCV DNA by PCR as described above. In all the transmission tests, *A. craccivora* Robinia was used as a positive control of ALCV transmission.

#### 2.3.2. EcmLV transmission

Aphid transmission of EcmLV was tested with *A. gossypii*, *M. persicae*, *D. plantaginea* and *A. craccivora* EuphorbiaSA (Table 1). The virus acquisition feeding was performed on tomato, buckhorn plantains, and euphorbia medusa’s head plants agroinfected with EcmLV. Plants used for virus inoculation by aphids were 2-week old tomato plants, one-month-old buckhorn plantain plants, or 6-year old euphorbia medusa’s head plants. The transmission success was assessed by PCR-mediated detection of EcmLV DNA. All the transmission tests of EcmLV were carried out in a P3 containment chamber.

#### 2.3.3. PlLV transmission

Aphid transmission of PlLV was tested with *D. plantaginea* and *A. gossypii*. Aphids were given access to PlLV by rearing them on buckhorn plantain plants one month after their agroinoculation with PlLV. Aphids thus exposed to PlLV, were shifted onto 1-month-old buckhorn plantain plants. The transmission success was assessed by PCR-mediated detection of PlLV DNA as described above.

### 2.4. Virus persistence and localization in aphids

#### 2.4.1. Testing virus persistence

Adults of *A. pisum*, *A. craccivora* Robinia and *A. craccivora* EuphorbiaSA were given a 3-day period on healthy broad bean plants for larvae delivery. Groups of 50 L1-L2 larvae were given a 3-day AAP on broad bean plants one month after their agroinoculation with ALCV. The growth conditions were 14h light at 26±2°C, and 10h dark at 24±2 °C. L3-L4 larvae were then moved on about 15 eight-day-old healthy plants - 10 individuals per plant-, and allowed for an IAP period of 2 days. The same individuals were shifted two more times to healthy plants. Individuals were sampled before AAP, after AAP and at the end of the third IAP, i.e. 6 days after the end of the AAP.

#### 2.4.2. Sampling of insect material

After collection, aphids were stored at -20°C until use. Some aphids were dissected by pulling the insect’s head with forceps under a binocular microscope. Gut and heads were separated in a water bath to prevent contaminations and subsequently grouped by 10 in 100 µL Edwards buffer. One drop of hemolymph was collected from each aphid with a glass capillary after pulling a leg. The DNA extraction was performed by grinding 10 individuals or organs with 30 rotations of small pestles whose conical tip match the internal volume of the conical shaped tip of 1.5 mL microtubes used as mortar. The crude extracts were centrifuged at 4000 g for 5 min. Supernatants were transferred on the filter of 200 µL filtered-tips with a PCR plate below and then centrifuged 10 min at 1500 g. One volume of isopropanol was added to the filtered extract. After several tubes inversions, the mix was centrifuged 25 min at 5000 g. Pellets were resuspended with 70% EtOH, and after centrifugation at 15700 g for 10 min, pellets were dried at 60°C and resuspended in 50µL H_2_O. DNA extracts were stored at -20°C before use.

#### 2.4.3. qPCR conditions

Amplification was performed with the LightCycler FastStart DNA Master Plus SYBR Green I kit (Roche) and the LightCycler 480 thermocycler (Roche). Primers were those described above for ALCV detection by PCR (ALCV2cEcmLV-F & ALCV2cEcmLV-R). They were used at a final concentration of 0.6 µM. Viral DNA detection was done by the addition of a 2 µL volume of extracted DNA to each well containing the Master-mix. The cycling protocol was as follows: an initial cycle consisting of 10 min at 95°C, 30 sec at 60°C and 20 sec at 72°C; 40 cycles consisting of 15 sec at 95°C, 30 sec at 60°C and 20 sec at 72°C; and finally a melting curve. DNA accumulations were reported with fluorescence values adjusted for amplification efficiencies with the LinRegPCR program (Ruijter et al., 2013). They were also presented as copy numbers of viral DNA with standard curves derived from 10 fold serial dilutions of recombinant plasmids containing the viral genome.

#### 2.4.4. Fluorescent in situ hybridization (FISH)

A fluorescent probe complementary to the CP gene of ALCV was prepared by random priming with the BioPrime DNA labeling system (Invitrogen) and Alexa Fluor 488-labeled dUTP. The template DNA was PCR amplified from the recombinant plasmid containing the ALCV genome, with the following primer pair: ALCV_FISH_620-F, 5’-GAA GAG GGC GAG AAC GAC AG-3’ and ALCV_FISH_1025-R, 5’-GTG GTC TAT TTC AGC AGT TGC C -3’. Just before use, 10 µL probe was diluted with 290 µL hybridization buffer (see below), denatured 10 min at 100°C and rapidly cooled on ice for 15 min. Individuals of *A. pisum*, *A. craccivora* Robinia and *A. craccivora* EuphorbiaSA were each divided into two groups raised for 2 weeks either on broad bean plants agroinfected with ALCV or on non-infected plants. Three days before use, aphids were shifted to healthy plants. Individuals were dissected in 1x phosphate-buffered saline (PBS) under a stereomicroscope. Pairs of salivary glands and digestive tracts were detached from the whole body by pulling the head with forceps. The dissected organs were fixed for 20 min at room temperature (RT) in embryo dishes containing 4% paraformaldehyde (PFA) diluted in PBS. Fixation was stopped by a 15-minute incubation in PBS containing 0,1M glycine. To improve the permeability of the tissues, they were incubated in H_2_O_2_ for 15 min. The dissected organs were then soaked 3 times 5 min in 20 mM Tris-HCl hybridization buffer (pH8) containing 0.9M NaCl, 0.01% SDS and 30% formamide. Organs were then incubated overnight at 37°C in the diluted and heat-denatured probe solutions (see above) in embryo dishes sealed with parafilm membranes. After three washing steps of 5 min with hybridization buffer and two with PBS, organs were mounted on microscope slides in Vectashield antifade mounting medium containing 1.5µg/mL DAPI for staining nuclei. Observations were performed using a Zeiss Confocal microscope and acquired in a stack mode. A minimum of thirty midguts and pairs of primary salivary glands were observed for ALCV exposed aphids of the three populations. Non-exposed aphids were observed as negative controls. The size of ALCV aggregates observed in midguts and salivary glands was estimated with the ImageJ software. To do this, the areas of 250 virtual confocal sections of aggregates were measured in midguts and salivary glands of 3 viruliferous individuals of *A. craccivora* Robinia and in the midgut of 3 viruliferous individuals of *A. craccivora* EuphorbiaSA.

### 2.5. Statistical analysis

All statistical analyses were conducted with the R software v3.6.1 (R Core Team, 2017). Data were compared using the Kruskal–Wallis rank sum test (function *krustal.test* of the package *stats*). When the null hypothesis of mean equality was rejected, the means of each pair of modalities were compared using the multiple comparison method based on the Benjamini and Yekutieli, 2001 procedure (function *pairwise.t.test()* with p-value adjustment method: BY).

## 3. Results

### 3.1. Aphid transmission is a common feature of capulaviruses

Aphid transmission of PlLV, previously shown with a non-cloned virus (Susi et al., 2017) and a Finnish *D. plantaginea* population (Susi et al., 2019), was tested here with an agroinfectious clone and a French population of *D. plantaginea*. Fifty percent of the Buckhorn plantain plants agroinoculated with PlLV were PCR positive, and as expected from the latent status of this virus (Susi 2017, 2019), none of them exhibited any particular symptom. Some of the agroinfected plants were used as source plants for a transmission test (Suppl. Table S1). The transmission was successful for nine of the ten test plants (Table 1) confirming that *D. plantaginea* is a vector of PlLV, and together with Susi et al. results, showed that transmission was possible irrespective of the geographic origin of the aphids, Finland or France.

The favored aphid candidate for the potential transmission of EcmLV was a black-backed aphid from South Africa collected on plants of the species *E. caput-medusae*, the natural host of EcmLV. These aphids were able to transmit EcmLV from an agroinoculated source plant to two *E. caput-medusae* plants obtained from seeds in controlled conditions (Table 1).

### 3.2. The aphid vector of EcmLV belong to the A. craccivora group

According to the sequence of the COI gene, individuals of the rearing population derived from black-backed aphids collected on *E. caput-medusae* in South-Africa clustered with representatives of the *A. craccivora* group, including *Aphis tirucallis* Hille Ris Lambers, 1954, a species known in Africa on *Euphorbia spp.* plants (Fig. 1). There are numerous morphologically similar *Aphis* spp. on *Euphorbia* that look like *A. craccivora* and which are recognized to be very difficult to distinguish from each other (Blackman and Eastop, 2000). As shown by morphological measures of specimens mounted on slides (Supplemental Fig.S1), the black-backed aphid specimens collected on Euphorbia medusa’s head plants are distinguishable from individuals of the *A. craccivora* group, *sensu stricto*, particularly by siphunculi length and a contrasting length ratio between the processus terminalis and the base of the last antennal segment (Supplemental Fig.S1), two previously reported discriminating features between specimens of the *A. craccivora* group, *sensu stricto*, and specimens of other *Aphis* living on *Euphorbia*, in particular, *Aphis euphorbiae* Kaltenbach, 1843 (Blackman and Eastop, 2000). Morphometric data suggest that our specimens probably belong to *A. tirucallis* or *A. euphorbiae species* but with low certainty because it cannot be excluded that the morphological differences are adaptive. Thus, as suggested by Coeur d’Acier et al. (2014) in this case, we adopted a pragmatic and conservative point of view and considered that it is safe to identify the black-backed aphids from South Africa as members of the *A. craccivora* group according to CO1 sequences. Hence, the name EuphorbiaSA was given to these South African aphids which refer both to their host plant and their geographical origin. Their adaptation to spurges was confirmed with branches cut from *Euphorbia nicaeensis* All. and *Euphorbia serrata* L. plants of the area of Montpellier, on which they developed readily.

**Figure 1.**
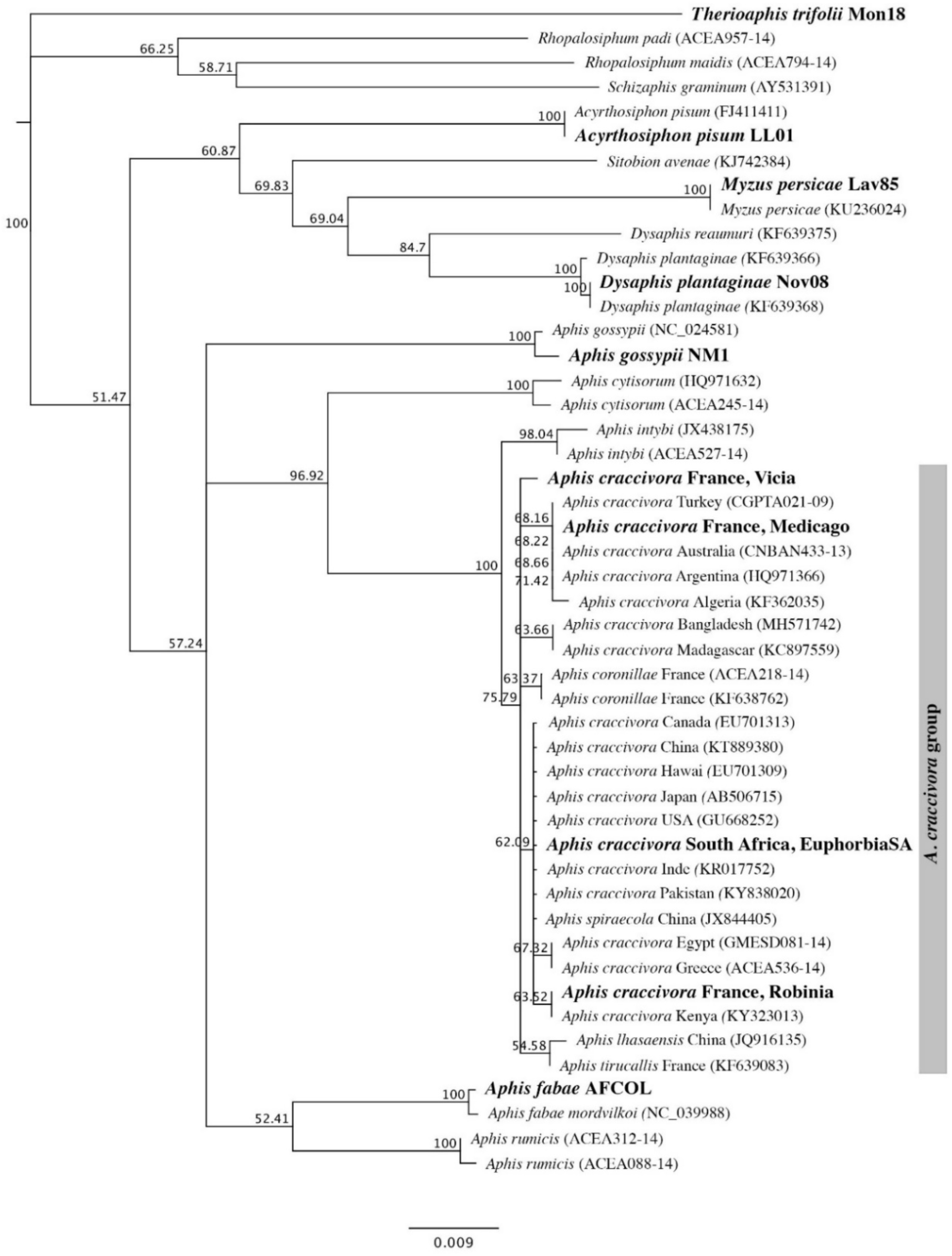
Neighbor joining tree showing the genetic distances among members of the *Aphis craccivora* group and members of related species, based on the cytochrome oxidase 1 gene. Sequences are identified by a species name and an accession number for those uploaded from Genbank or BOLD. Members of the *A. craccivora* group are identified further with the country of origin. The description of sequences generated in this study is in bold. A detailed description of the sequenced specimens is provided in Supplemental Tables S1 and S2. Numbers associated with nodes represent the percentage of 10,000 bootstrap iterations supporting the nodes; only percentages > 50 % are indicated. *Therioaphis trifolii* was used as an outgroup.

### 3.3. The transmission of capulaviruses by aphids is highly specific

Aphids of several species were tested for their ability to transmit ALCV (Table 1). They were selected according to at least one of the following criteria: regularly found on Fabaceae, phylogenetically close to the known vectors of capulaviruses, and reported to transmit other viruses. Although aphids of the species *A. fabae* and *A. gossypii* belong to the same genus as the vector aphids *A. craccivora*, they did not transmit ALCV. *Myzus persicae* and *A. pisum* are polyphagous species that were both reported to transmit circulative persistent viruses. However, they were not detected as vectors of ALCV. *Therioaphis trifolii*, is a specialist of alfalfa, the species on which ALCV was isolated and on which it was mostly reported (Davoodi et al., 2018). However, it was not detected as a vector of ALCV. The transmission tests conducted with EcmLV and PlLV were restricted by the limited number of aphids available in the laboratory that develops on the few reported hosts of these viruses, i.e. *A. gossypii* and *M. persicae*. EcmLV was not transmissible by *A. gossypii* and *M. persicae* and PlLV was not transmissible by *A. gossypii* (Table 1). These transmission failures, with the detection of only one vector species per capulavirus species, are consistent with a relatively high transmission specificity.

Transmission specificity was explored further by testing if an aphid vector of one capulavirus can be a vector of another one. Due to a limited number of compatible host combinations of virus and aphid species (Table 3A), such transmission specificity could be tested thoroughly with one combination only, i.e *A. craccivora* EuphorbiaSA and ALCV which are both hosted by the broad bean. According to symptom observations and the ALCV specific PCR test, all the broad bean plants exposed to EuphorbiaSA aphids that were given access to ALCV infected plants, were negative for the presence of ALCV (Table 1: 0/12, 0/45, 0/15). The transmission was also unsuccessful in test plants exposed to 30-150 viruliferous aphids (0/6). Aphids of the *A. craccivora* Robinia population tested in parallel as positive controls did successfully transmit ALCV (Table 1: 8/15, 9/30).

To explore other heterologous combinations of capulaviruses and known capulavirus vectors for transmission, we tested if capulaviruses available in the laboratory have common host plants on which transmission tests may be performed (Table 3B). Heterologous infection was possible only for one of 20 ALCV inoculated buckhorn plantain plants and one of 19 EcmLV inoculated buckhorn plantain plants. The ALCV infected buckhorn plantain tested PCR-negative two months after agroinoculation. Hence, as ALCV infection was transient, we only tested the transmission of EcmLV by *D. plantaginea*. The 12 buckhorn plantains plants used for the IAP were PCR-negative for the detection of EcmLV (Table 1). This result may suggest that EcmLV is not transmitted by *D. plantaginae* but it cannot be excluded that the infection failure was due to the fact that plantain is a poor host of EcmLV.

### 3.4. Transmission specificity of ALCV within the A. craccivora group

The results presented above show that while aphids of the Robinia population of *A. craccivora* were able to transmit ALCV, those of the EuphorbiaSA population, were not (Table 1). This indicates that the transmission of ALCV by aphids is highly specific. Hence, the specificity within the *A. craccivora* group was investigated further by testing two more populations, Medicago and Vicia (Supplemental Table S1). Aphids of Robinia, Vicia and Medicago populations transmitted ALCV from broad bean agroinfected plants to 7 of 13 (53.8%), 7 of 20 (35%) and 1 of 20 (5%) test plants (Table 2A). The relatively low transmission success with the Medicago population may be explained by its low affinity for the broad bean as observed during the transmission test in which some individuals did not stay on broad bean plants. This hypothesis is consistent with a higher transmission success observed in a preliminary transmission test with alfalfa. Indeed, aphids of the Medicago population transmitted ALCV to 2 of 4 alfalfa test plants, following an access period on a naturally ALCV infected alfalfa plant collected near Montpellier (Table 2A).

**Table 2.**
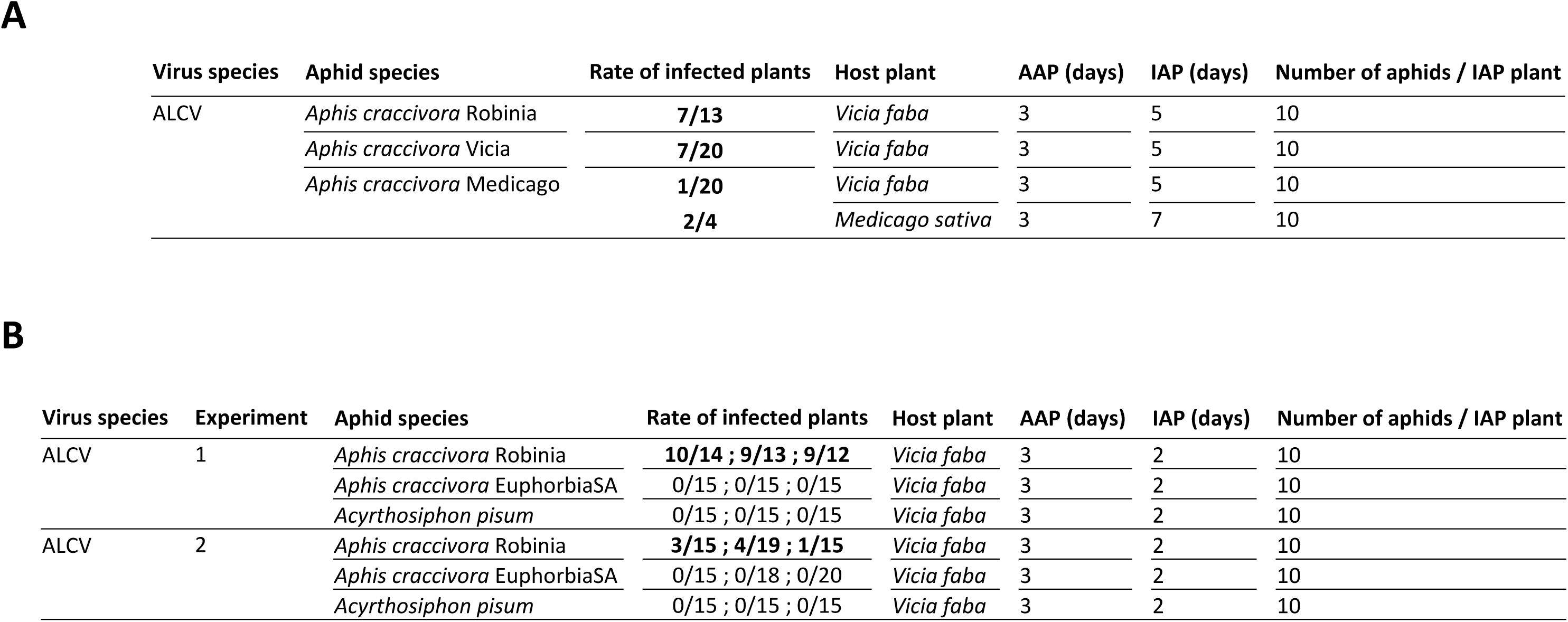
Transmission tests of ALCV with *A. craccivora* populations. (A) Comparison of populations from *Robinia pseudoacacia* L. (Robinia), *Vicia sativa* (L.) Bernh. (Vicia), and *Medicago sativa* L. (Medicago) (B) Test of the persistency of the infectivity in *A. craccivora* Robinia. The persistency was assessed with individuals that were given a 3-day AAP on broad bean plants agroinfected with ALCV and subsequently shifted sequentially to three batches of plants each exposed to 2-day IAPs. The rate of transmission successes is mentioned in the chronological order of the three passages. Aphids of the EuphorbiaSA population of *A. craccivora* and of *A. pisum* were used as negative controls particularly for the persistence of ALCV DNA determined in the whole body (Experiment 1, Fig. 2) and in dissected fractions (Experiment 2, Fig. 3).

**A**
| Virus species | Aphid species | Rate of infected plants | Host plant | AAP (days) | IAP (days) | Number of aphids / IAP plant |
| --- | --- | --- | --- | --- | --- | --- |
| ALCV | *Aphis craccivora* Robinia | **7/13** | *Vicia faba* | 3 | 5 | 10 |
|  | *Aphis craccivora* Vicia | **7/20** | *Vicia faba* | 3 | 5 | 10 |
|  | *Aphis craccivora* Medicago | **1/20** | *Vicia faba* | 3 | 5 | 10 |
|  |  | **2/4** | *Medicago sativa* | 3 | 7 | 10 |

**Table 2.** Transmission tests of ALCV with *A.
| Virus species | Experiment | Aphid species | Rate of infected plants | Host plant | AAP (days) | IAP (days) | Number of aphids / IAP plant |
| --- | --- | --- | --- | --- | --- | --- | --- |
| ALCV | 1 | *Aphis craccivora* Robinia | **10/14 ; 9/13 ; 9/12** | *Vicia faba* | 3 | 2 | 10 |
|  |  | *Aphis craccivora* EuphorbiaSA | 0/15 ; 0/15 ; 0/15 | *Vicia faba* | 3 | 2 | 10 |
|  |  | *Acyrtosiphon pisum* | 0/15 ; 0/15 ; 0/15 | *Vicia faba* | 3 | 2 | 10 |
| ALCV | 2 | *Aphis craccivora* Robinia | **3/15 ; 4/19 ; 1/15** | *Vicia faba* | 3 | 2 | 10 |
|  |  | *Aphis craccivora* EuphorbiaSA | 0/15 ; 0/18 ; 0/20 | *Vicia faba* | 3 | 2 | 10 |
|  |  | *Acyrtosiphon pisum* | 0/15 ; 0/15 ; 0/15 | *Vicia faba* | 3 | 2 | 10 |

### 3.5. ALCV persistence in vector and non-vector populations of the A. craccivora group

Geminiviruses are transmitted by insects in a circulative persistent manner (Nault, 1997, Whitfield et al., 2015). As capulaviruses were readily transmitted with long AAPs and IAPs (Roumagnac et al., 2015 and this study) it was supposed that they also transmitted in a circulative persistent manner. The success of this mode of transmission depends upon the ability of the virus to cross the gut and the salivary epithelia of the insect vectors and to avoid defense and degradation mechanisms. Considering the close relationship between aphids of the *A. craccivora* group, it was thought that non-vector aphids of the EuphorbiaSA population may reflect some of the features of vector aphids regarding ALCV transmission. This prediction was tested by assessing viral persistence in EuphorbiaSA and Robinia populations of *A. craccivora,* in comparison with *A. pisum* an out-group non-vector control. The transmission procedure consisted of a 3-day AAP on broad bean plants agroinfected with ALCV, followed by three sequential 2-day IAPs on three batches of broad bean plants. While 70% of the test plants of the Robinia treatment were symptomatic in the three batches, none of the test plants exposed to individuals of *A. pisum* and EuphorbiaSA populations exhibited symptoms, which confirmed their non-vector status (Table 2B). The infection in the three sequentially inoculated Robinia batches shows that ALCV persists in the Robinia aphids at least up to 4 days after their withdrawal from the viral source. To know to which extent ALCV may also persist in the non-vector aphids of *A. pisum* and *A. craccivora* EuphorbiaSA, viral DNA content was assessed by qPCR. At the end of the 3-day AAP, ALCV DNA was detected in vector and non-vector aphids showing that all had access to the virus (Fig. 2A). The number of viral DNA copies per insect in *A. craccivora* Robinia (1.5×10^6^) was significantly higher than that of *A. pisum* (1.6×10^5^) and *A. craccivora* EuphorbiaSA (5.6×10^4^) (Fig. 2A). At the end of the three IAPs, the number of viral copies estimated for *A. craccivora* Robinia was in average 7.5×10^5^ viral copies per insect, significantly higher than that of *A. craccivora* EuphorbiaSA (6.4×10^4^) (Fig. 2B). On the other hand, the number of copies in *A. pisum* aphids, if any, was below the detection threshold. These results show that after withdrawing aphids from the virus source, ALCV can persist at least up to 6 days in *A. craccivora* aphids, irrespective of the population, but not in *A. pisum* aphids in which ALCV was not detected at this time point.

**Figure 2.**
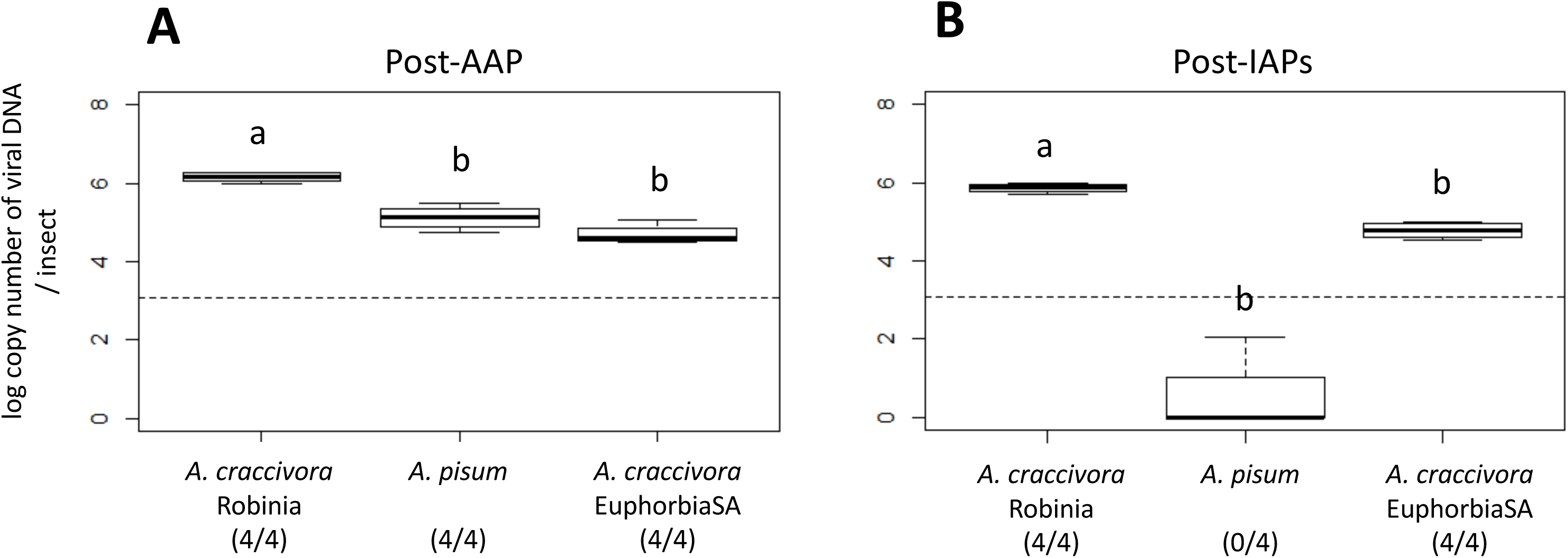
Box-plots showing the amount of ALCV DNA in vector aphids (*A. craccivora* Robinia) and non-vector aphids (*A. craccivora* EuphorbiaSA and *A. pisum*) following a 3-day AAP on broad bean plants agroinfected with ALCV (A), and after three sequential 2-day IAPs on non-infected plants (B). In each treatment, the content of ALCV DNA was determined by qPCR in 4 pools of 10 individuals. Accumulations are reported as logarithm 10 of the number of viral DNA copies. The dotted lines represent the highest values obtained with non-viruliferous pools of 10 aphids sampled before the 3-day AAP; the post-AAP and IAPs pools for which the number of estimated DNA copies is above this threshold are considered positive for the presence of ALCV DNA. The ratio of positive pools is indicated below aphid names. Letters above box-plots indicate significant differences (p-value < 0.01) between the modalities according to the multiple comparison method based on Benjamini and Yekutieli’s procedure.

Interestingly, the results observed with *A. craccivora* EuphorbiaSA with respect to ALCV/aphid interactions, were intermediate between those observed with *A. craccivora* Robinia and *A. pisum*. Indeed, like *A. pisum*, *A. craccivora* EuphorbiaSA was not a vector of ALCV, and like *A. craccivora* Robinia, it retained ALCV DNA. The hypothesis of 6-day retention of viral DNA in the gut lumen is quite unlikely because *A. pisum* aphids, whose body is about three times larger than that of EuphorbiaSA aphids, tested negative at this time point. Therefore, it was thought that after internalization in gut cells of EuphorbiaSA aphids, ALCV may avoid to some extent degradation pathways and possibly circulate further in the hemocoel and other organs like salivary glands. To test this hypothesis a second transmission test was carried out with the same procedure as the first test, except that the viral quantification was not done on the whole body, but on dissected parts, i.e. the digestive tract, the head with the salivary glands, and hemolymph. The three aphid parts were from 10 aphids in each qPCR sample. Like in the first test, symptomatic broad bean plants were observed only in the Robinia treatment (Table 2B). At the end of the AAP, viral DNA was detected in the digestive tract in all the treatments (Fig. 3A) which shows that aphids of the three populations had access to the virus. Although the body of *A. pisum* aphids is much larger than that of EuphorbiaSA aphids, the average ALCV DNA content per insect was similar (1.75×10^4^ and 1.4×10^4^, respectively). The guts of Robinia aphids contained 2.0×10^6^ viral DNA copies per insect on average which is 145 times more than the content estimated for *A. pisum* and EuphorbiaSA. At the end of the three sequential 2-day IAPs, the digestive tract of *A. pisum* aphids was qPCR negative for the detection of ALCV (Fig. 3B). On the contrary, the samples of digestive tracts of Robinia aphids were qPCR positive (6.7×10^4^ viral DNA copies per individual) as well as 4 of the 6 samples of EuphorbiaSA (5.7×10^2^ viral DNA copies per individual). Only the hemolymph samples of Robinia aphids were detected qPCR-positive for the detection of ALCV, with 5.1×10^2^ viral DNA copies per individual. Finally, the head samples were qPCR negative for *A. pisum*, positive for Robinia with 5.8×10^3^ viral DNA copies per individual, and partially positive for EuphorbiaSA, with 5 of 7 samples being positive (1.9×10^2^ viral DNA copies per individual).

**Figure 3.**
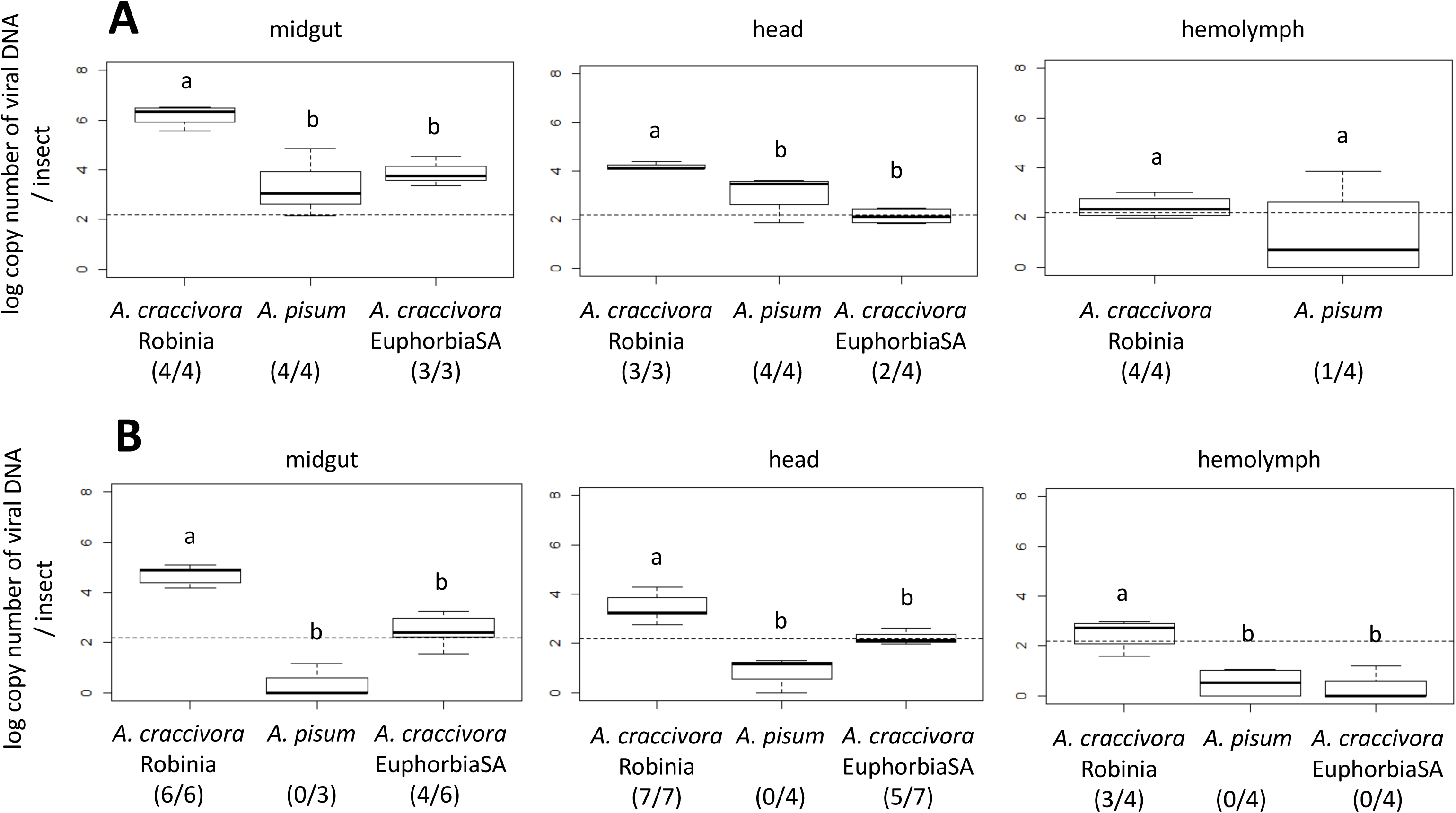
Box-plots showing amount of ALCV DNA in the midgut, head, and hemolymph in vector aphids (*A. craccivora* Robinia) and non-vector aphids (*A. craccivora* EuphorbiaSA and *A. pisum*), following 3-day AAP on broad bean plants agroinfected with ALCV (A) and after three sequential 2-day passages on non-infected plants (B). The content of ALCV DNA was determined by qPCR in 3-7 pools of 10 dissected fractions. Accumulations are reported as logarithm 10 of the number of viral DNA copies. The dotted lines represent the highest value obtained with non-viruliferous pools of 10 aphids sampled before the 3-day AAP; the post-AAP and IAPs pools for which the number of estimated DNA copies is above this threshold are considered positive for the presence of ALCV DNA. The ratio of positive pools is indicated below the aphid names. Presentation of accumulations and statistical analysis as in Fig. 2.

### 3.6. Intracellular ALCV aggregates were localized in vector and non-vector A. craccivora

To confirm that the persistence of ALCV detected with the non-vector population of *A. craccivora* (EuphorbiaSA) may be associated with internalization and persistence in midgut cells, we used fluorescent in-situ hybridization (FISH) of ALCV DNA. *Aphis craccivora* Robinia and *A. pisum* were used as positive and negative controls, respectively. Aphids of the three populations were allowed 15-day AAPs followed by 48-hour IAPs before dissection and FISH analysis. In Robinia aphids, fluorescent aggregates were observed all around the nuclei of epithelial cells of the anterior midgut (Fig. 4) and of the beginning of the posterior midgut (data not shown). Their area estimated from confocal virtual sections was 0.52 ± 0.02 µm^2^ on average (Fig. 6). These aggregates were detected neither in aphids of the *A. pisum* population nor in aphids that fed on healthy plants only. In EuphorbiaSA aphids, virus-specific aggregates were also detected but were distinguishable from those observed with Robinia aphids by their distribution and size. Indeed, unlike those of Robinia aphids, they did not circle the nuclei but seemed to be concentrated towards the apical and basal side of epithelial cells. Their area was 28.10 ± 1.01 µm^2^ on average, significantly larger than that of the Robinia aggregates (Fig. 6, Benjamini and Yekutieli’s method: p-value < 0.01). It is noteworthy that the distribution of aggregates’ sizes is an additional distinguishing feature between the two *A. craccivora* populations. Additionally, while virus aggregates were observed in a majority of Robinia individuals (68%), the aggregates were observed in only 33% of EuphorbiaSA individuals (Table 4).

**Figure 4.**
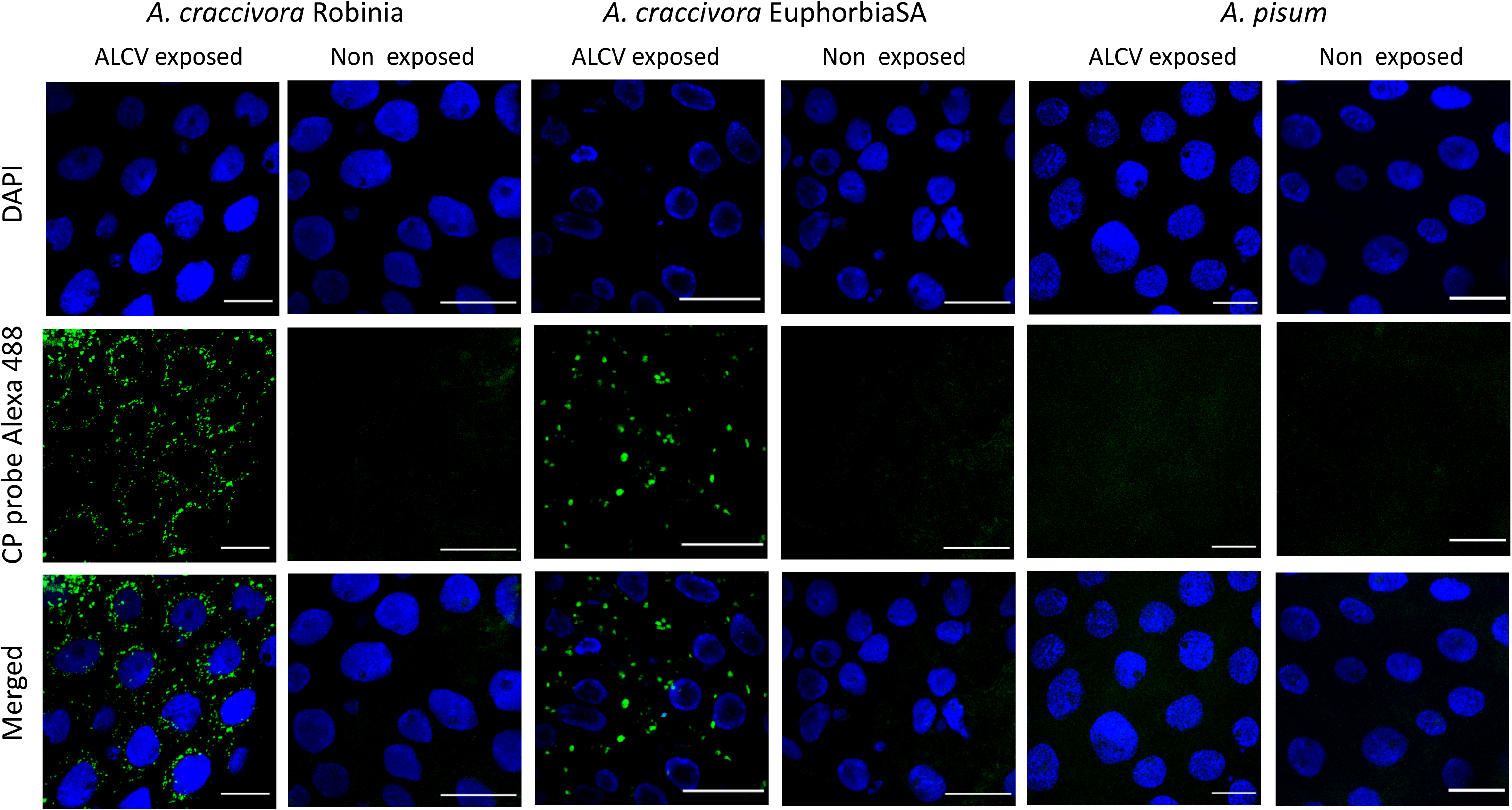
Localization of ALCV DNA by FISH in dissected anterior midguts of vector and non-vector aphids of ALCV. Aphids were exposed during 15 days to broad bean plants agroinfected with ALCV and then shifted to non-infected plants for three days before FISH analysis. Non exposed aphids underwent the same procedure except that broad bean plants of the 15 day period were non-infected. *A. craccivora* Robinia is a vector aphid of ALCV whereas *A. craccivora* EuphorbiaSA and *A. pisum* are non-vector aphids. The ALCV specific DNA probe is labeled with a green Alexa 488 fluorochrome. Nuclei are DAPI-blue stained. Preparations were examined with confocal microscopy. Horizontal bars = 30 µm.

**Figure 5.**
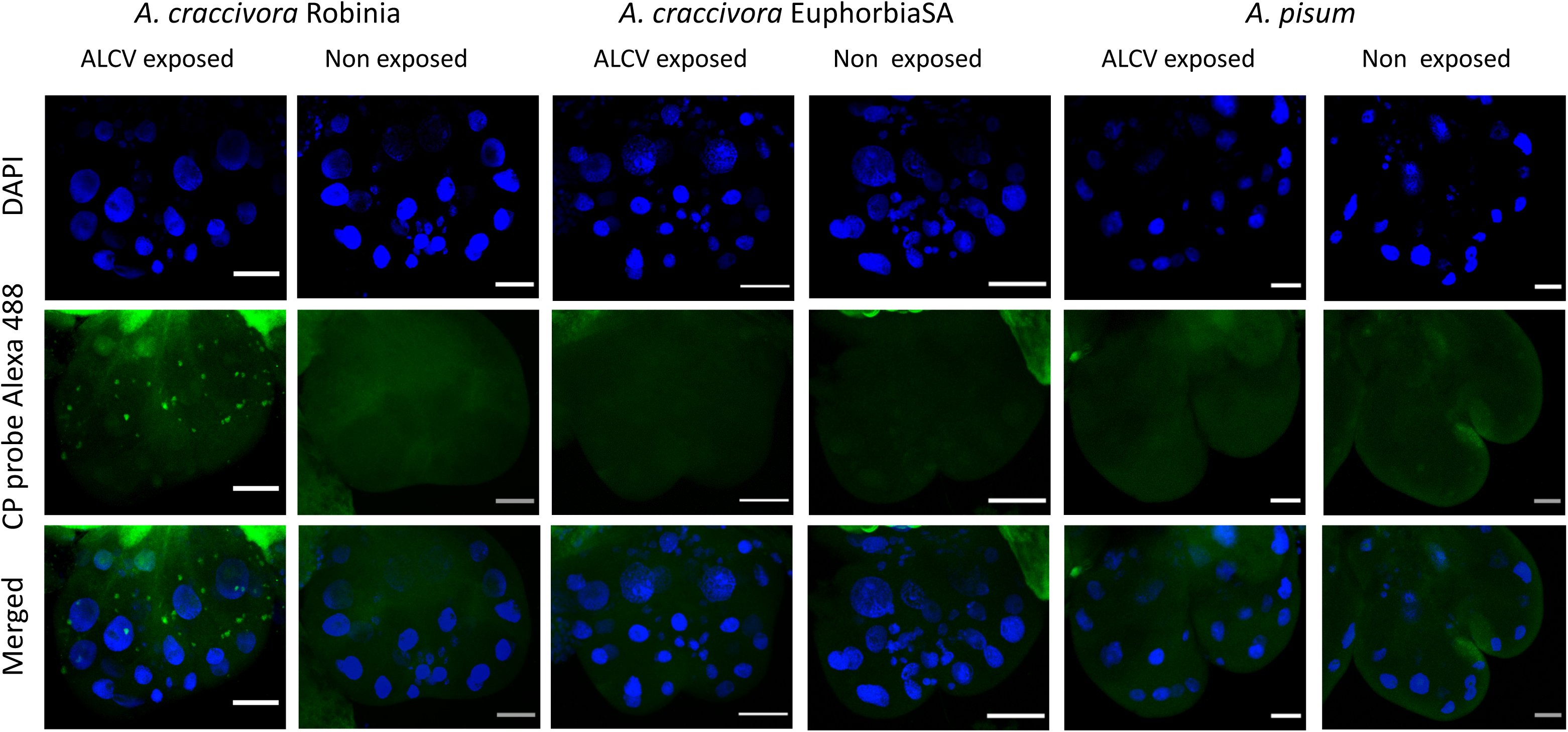
Localization of ALCV DNA by FISH in dissected primary salivary glands of vector and non-vector aphids of ALCV. Aphid species, figure layout, probe and DAPI staining as in Fig. 4. Horizontal bars = 30 µm.

**Figure 6.**
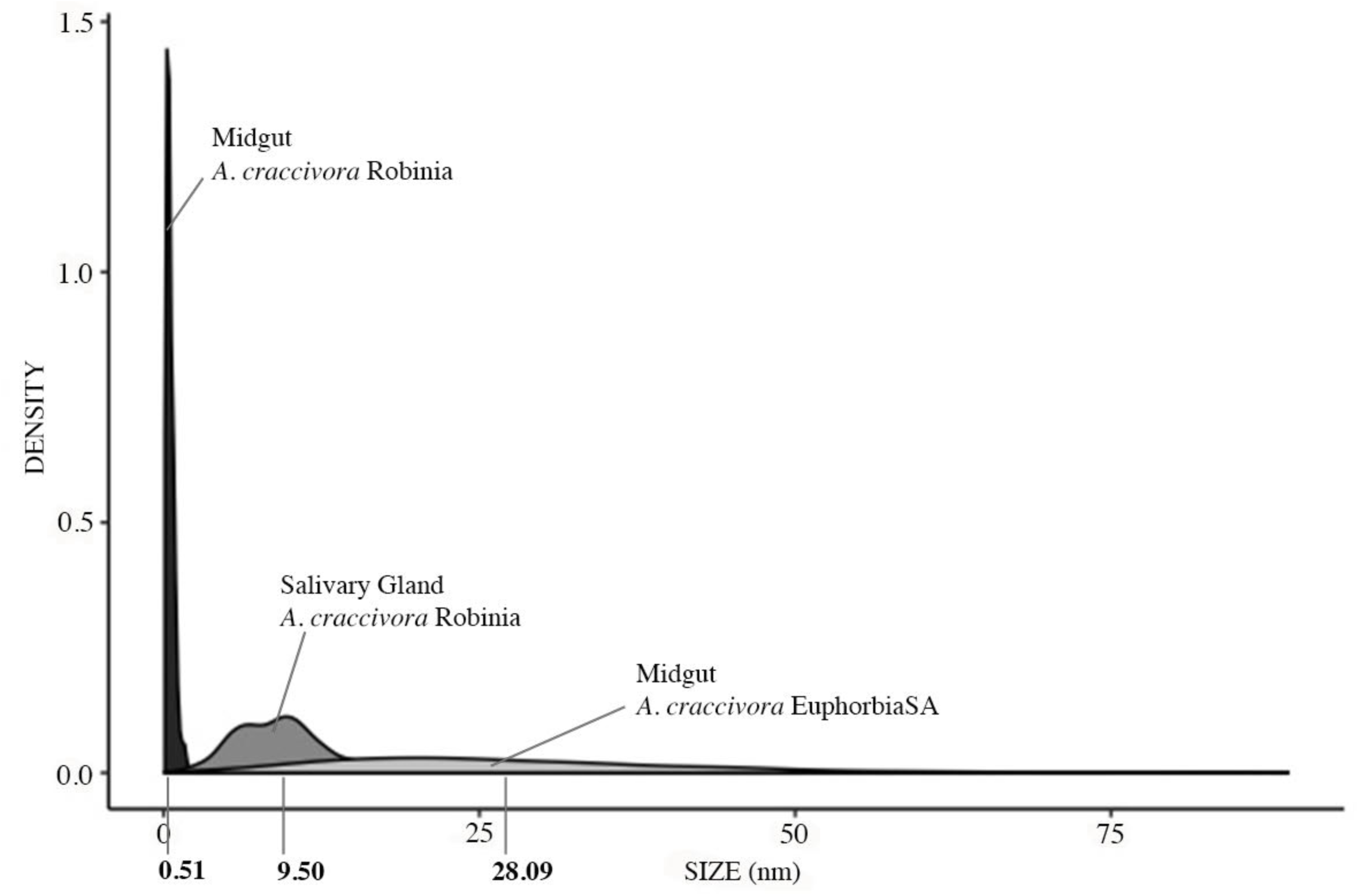
Density plots showing the frequency distribution of section areas of fluorescent ALCV-specific aggregates observed in dissected midguts and salivary glands of vector and non-vector *A. craccivora* aphids. FISH was performed on aphids that were given a 15-day AAP on broad bean plants agroinfected with ALCV, followed by a 3-day period on healthy plants. Areas of aggregate sections were measured in midguts and primary salivary glands of three vector aphids (*A. craccivora* Robinia) and in midguts of three non-vector aphids (*A. craccivora* Euphorbia SA). Areas of 250 sections of aggregates were measured per modality. A graph was obtained with the libraries *ggplot2* and function *ggplot* in R.

**Table 3.**
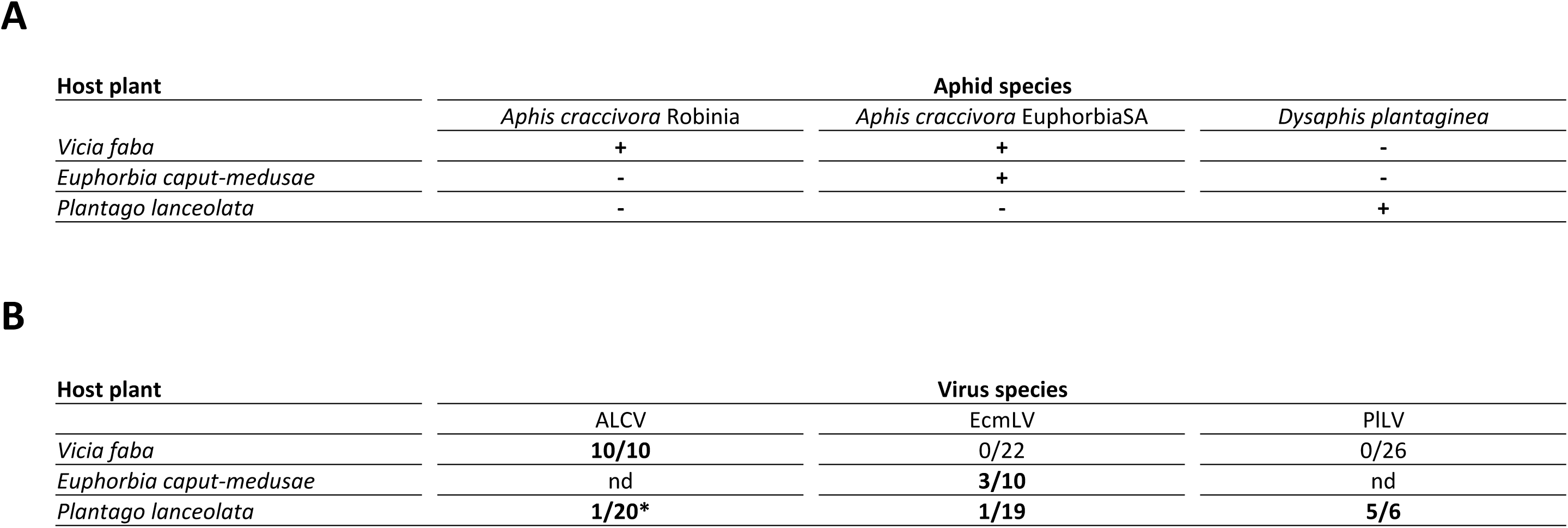
Detection of compatible plant/aphid/capulavirus trios to assess the transmission specificity of capulaviruses by aphids. (A) reported host plants of capulaviruses that may be hosts of capulavirus vectors. + aphids were able to stay for at least several days on the plant; - aphids were not able to survive on the plant. (B) Detection of reported host plants of capulaviruses that may host other capulaviruses. Capulaviruses were agroinoculated to host plants. nd = not done : * = transitory virus infection.

**A**
| Host plant | Aphid species |  |  |
| --- | --- | --- | --- |
|  | *Aphis craccivora* Robinia | *Aphis craccivora* EuphorbiaSA | *Dysaphis plantaginea* |
| *Vicia faba* | + | + | - |
| *Euphorbia caput-medusae* | - | + | - |
| *Plantago lanceolata* | - | - | + |

**Table 3.** Detection of compatible plant/aphid/capulavirus trios to assess the transmission specificity of capulaviruses by aphids.
| Host plant | Virus species |  |  |
| --- | --- | --- | --- |
|  | ALCV | EcmlV | PILV |
| *Vicia faba* | **10/10** | 0/22 | 0/26 |
| *Euphorbia caput-medusae* | nd | **3/10** | nd |
| *Plantago lanceolata* | **1/20\*** | **1/19** | **5/6** |

**Table 4.**
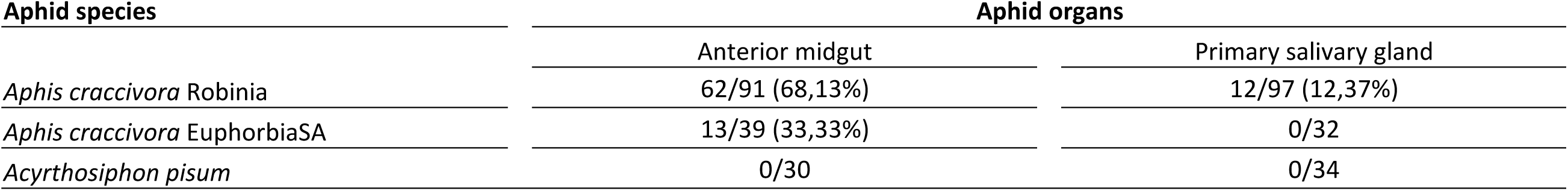
Frequency of midgut and salivary glands in which ALCV-specific aggregates were detected by FISH in vector and non-vector aphids (Figs 2 and 3). Vector aphids were from the Robinia population of *A. craccivora* and the non-vector aphids were from the EuphorbiaSA population of *A. craccivora* and from *A. pisum*. Aphids were analyzed after a 15-day AAP on broad bean plants agroinfected with ALCV followed by a 3-day IAP on non-infected plants.

Virus-specific aggregates were detected in primary salivary glands of *A. craccivora* Robinia aphids and only in 13% of individuals (Table 4). Interestingly, their size was larger than the Robinia midgut aggregates, with a mean area of 9.50 ± 0.62 µm^2^ (Fig. 6). No virus-specific aggregates were detected in salivary glands of *A. craccivora* EuphorbiaSA and *A. pisum* aphids.

## 4. Discussion

### 4.1. Aphid transmission is a taxonomic criterion of the genus Capulavirus

The insect vector is an important criterion for defining genera within plant virus families (Zerbini et al., 2017). Aphid transmission was reported for ALCV and PlLV, two of the four members of the Capulavirus genus. It suggested that aphid transmission may be a distinctive feature of the genus. By demonstrating that a third capulavirus (EcmLV) is transmitted by aphid as well, aphid transmission is validated as a taxonomic criterion of the new genus. Additionally, the successful transmission of a cloned PlLV by the aphid *D. plantaginea* not only confirmed previous transmission results with a non-cloned PlLV isolate (Susi et al., 2019), but showed, together with the transmission of ALCV (Roumagnac et al., 2015) and EcmLV (this study) clones, that their monopartite genome determines by itself aphid transmission without any helper DNA or virus. It is supposed that the CP of capulaviruses is a major player in aphid transmission because CP was reported with other geminiviruses to be required for transmission (Azzam et al., 1994) and insect specificity (Briddon et al., 1990). Consistently with these reports, it is noteworthy that the CPs of ALCV and EcmLV, the two capulaviruses transmitted by aphids of the *A. craccivora* group, are more similar to each other (75% amino acid identity) than to the CP of PlLV (47% and 53% aa identity respectively) which is transmitted with a non-craccivora aphid, i.e. *D. plantaginea*. Hence, it may be predicted that the fourth reported FbSLCV would also be transmitted with an *A. craccivora* aphid because its CP is highly similar to ALCV and EcmLV CPs (78% and 72%, respectively) (Varsani et al., 2017).

### 4.2. Transmission results and taxonomy of A. craccivora populations

The transmission tests showed that the *A. craccivora* group comprises two capulavirus vectors, the EuphorbiaSA population, vector of EcmLV, and three other populations (Robinia, Medicago, and Vicia), vectors of ALCV. Our molecular analysis with the barcoding gene COI (Fig. 1) confirms previous work showing the difficulty to differentiate sub-species/populations within the very cosmopolitan *A. craccivora* group (Blackman and Eastop, 2000, Wang et al., 2011; Song et al., 2016; Coeur d’Acier et al., 2014). The EuphorbiaSA population was distinguished from other *A. craccivora* populations by its adaptation to spurges and by morphological differences. However, according to aphid literature, it cannot be excluded that morphological variations are non-heritable adaptive features (Mehrparvar, 2012). Therefore, as recommended by these authors, morphological differentiation among populations needs to be confirmed with genetic and biological data to exactly evaluate their taxonomic situations. Unexpectedly, the contrasted interactions of ALCV with the *A. craccivora* populations may be a complementary biological feature to distinguish the EuphorbiaSA population from the other three *A. craccivora* populations used in this study. The soundness of such a distinguishing feature is supported by the transmission results of luteoviruses by aphids. Indeed, the intra-species transmission specificity previously detected with various populations of *Schizaphis graminum* (Rondani, 1852), an aphid vector of luteoviruses (Gray et al., 2002), was shown to be a heritable trait regulated by multiple genes acting in an additive fashion (Burrows et al., 2007).

### 4.3. Contrasted vector ranges among geminivirus genus

EcmLV was identified in South Africa and ALCV in Europe, Argentina and the Middle East (Davoodi et al., 2018) which suggest that *A. craccivora* is a global vector of capulaviruses. This prediction is consistent with the global distribution of *A. craccivora* and with the fact that FbSLCV, the Indian capulavirus, was isolated from French bean, a highly conducive host of *A. craccivora*. Thus, without PlLV, the transmission of capulaviruses may be comparable to the transmission of begomoviruses, both having a unique vector species, respectively, *A. craccivora* and the whitefly, *Bemisia tabaci* Gennadius, 1889. Nevertheless, unlike *A. craccivora* in which a population (EuphorbiaSA) was detected to be non-vector of an *A. craccivora* transmitted capulavirus, it can be inferred from the literature that *B. tabaci* populations are vector of any begomovirus, provided that the plant used for transmission is host of both the virus and the whitefly population (Burban et al., 1992). When the capulavirus genus is considered as a whole, including PlLV, the transmission of capulaviruses is comparable to the transmission of mastreviruses with vectors belonging to different genera (Kvarnheden et al., 2016).

Contrary to *A. craccivora*, *D. plantaginea* is a very specialist aphid that moves back and forth between buckhorn plantain and apple tree, its primary host. The contrasting breadth of the host ranges of capulavirus vectors is expected to have a direct effect on the natural host range of capulaviruses. Thus, it is expected that the host range of PlLV will be limited to apple trees. As far as we know, PlLV is the first described plant virus transmitted by *D. plantaginea*, contrary to *A. craccivora* which is known to transmit plant viruses belonging to many families, ie. *Tombusviridae*, *Potyviridae*, *Bromoviridae*, *Nanoviridae*.

### 4.4. ALCV is transmitted by A. craccivora in a highly specific and persistent circulative manner

The transmission of capulaviruses by aphids was supposed to be according to the circulative persistent mode. However, it had not been demonstrated so far. Here, as ALCV DNA could be detected in *A. craccivora* individuals 6 days post-AAP, and infectivity up to 4 days post-AAP, we demonstrate that the mode of transmission of ALCV is persistent. Moreover, the localization of ALCV DNA in epithelial cells of the midgut and the primary salivary glands support a circulative mode of transmission.

ALCV was not transmissible by the EuphorbiaSA population of *A. craccivora*. The transmission tests with a high number of viruliferous EuphorbiaSA individuals per test plants (30-150) excluded the possibility of a potentially low amount of inoculated virus due to the smaller size of EuphorbiaSA aphids (Supplemental Fig. S1). Thus, the failure of ALCV by *A. craccivora* EuphorbiaSA is expected to be due to incompatible virus-aphid interactions rather than to a potentially insufficient amount of inoculated virus. The extremely high transmission specificity of ALCV by certain populations of *A. craccivora* is similar to that reported with barley yellow dwarf virus - MAV (BYDV-MAV) and BYDV-PAV differentially transmitted by *S. graminum* populations (Gray et al., 2002). However, it was not detected in other geminivirus genera, not even within *B. tabaci* populations that are thought to be vector of all begomoviruses. The molecular interactions that drive ALCV recognition for transmission are expected to be highly specific. Hence, it seems highly unlikely that capulaviruses would be transmitted by non-aphid vectors, which in turn confirm that aphid transmission is a taxonomic criterion of the genus Capulavirus.

The incapability of the EuphorbiaSA population of *A. craccivora* to transmit ALCV may be related to its specialization in the transmission of EcmLV. It would be interesting to validate this hypothesis by a symmetrical test in which the transmission of EcmLV would be tested with *A. craccivora* populations that are vectors of ALCV. Unfortunately besides EuphorbiaSA, *A. craccivora* populations available in our laboratory did not develop on reported hosts of EcmLV (Bernardo et al., 2013). However, this complementary test may be carried out in the future with African *A. craccivora* populations reported from spurges (Remaudière, 1985).

### 4.5. Transmission barriers to ALCV in aphids

The detection of ALCV DNA in some hemolymph samples of the non-vector *A. pisum* species at the end of the AAP may be consistent with a permissive gut barrier. However, as most of the hemolymph samples of *A. pisum* were negative and ALCV was not localized in midgut epithelial cells by FISH, the positive detection in hemolymph may rather be contamination. Thus, the results obtained with the non-vector *A. pisum* are consistent with a transmission barrier located at the gut level similarly to the barriers detected with non-vector leafhoppers of mastreviruses (Lett et al., 2002) and non vector whiteflies of begomoviruses (Rosell et al., 1999; Czosnek et al., 2001; Ohnishi et al., 2009). The viral persistence in aphids of the non-vector EuphorbiaSA population of *A. craccivora* but not of *A. pisum* may be explained by the access of ALCV to the gut cells of EuphorbiaSA aphids but not to those of *A. pisum* aphids.

The results on ALCV persistence and localization in the body of the non-vector EuphorbiaSA population of *A. craccivora* suggest that the transmission barrier is in the salivary glands. Indeed, viral DNA was detected by FISH in midgut epithelial cells, and at 6 days post-AAP, some head samples were detected virus-positive. Interestingly, salivary glands were previously reported to be the major barrier for the aphid-transmitted barley yellow dwarf disease associated viruses (*Luteoviridae*) (Gildow and Gray, 1993). Indeed, these authors showed that the basal lamina (BL) surrounding the accessory salivary glands act as a viral barrier. It was observed to be completely tight to BYDV-MAV in the non-vector aphid *Rhopalosiphum maidis* (Fitch, 1856), slightly permissive in the non-efficient vector *Rhopalosiphum padi* Linnaeus, 1758 but attractive in the efficient vector *Sitobion avenae* (Fabricius, 1775).

ALCV DNA was readily localized by FISH in the midgut of vector aphids of *A. craccivora* species. Its rare observation in the salivary glands may be explained by difficult access to salivary glands due to the BL barrier as reported for BYDV-MAV in *R. maidis* (Gildow and Gray, 1993). Indeed, as these authors estimated the size exclusion of BL in cereal aphids to 20 to 30 nm (Peiffer et al., 1997), a potentially similar size exclusion in *A. craccivora* may reduce the capulavirus transit through the BL due to their size, 20 x 36 nm in the case of EcmLV (Roumagnac et al., 2015).

The standard size of the virus aggregates detected by FISH in midgut cells of the Robinia population individuals of *A. craccivora* and their defined perinuclear distribution reflect a well-established mechanism associated with virus transit (Fig. 6). On the contrary, the large range of sizes of the aggregates observed in midgut cells of EuphorbiaSA individuals and their apparently undefined distribution seems to reflect non established virus-aphid interactions which nevertheless allow some virus persistence. Further tests will be necessary to characterize the specific and non-specific interactions and confirm the transmission barriers of capulaviruses.

## Acknowledgements

We are very thankful to Armelle Coeur d’Acier for her valuable expertise in aphid identification. We thank Myriam Siegwaert (Inra-Avignon) for providing *D. plantaginea* aphids, Sylvaine Boissinot (Inra-Colmar) and Christoph Vorburger (ETHzürich) for providing *A. fabae* aphids and Josep Vicens Fandos (Lab. de Botánica, Fac. de Farmacia, Univ. de Barcelona) for confirming the species identification of the spurges collected near Montpellier. The design of the CO1 primers was from Isabelle Meunier (Inra-Montpellier). The study was carried out during the Ph.D. project of Faustine Ryckebusch that was funded by the Agropolis Fondation (E-Space flagship program) grant number 1504-004.

---

**Supplemental Figure 1.** Morphological comparisons between specimens of a population of *A. craccivora sensu stricto* (a, a’) and specimens of a euphorbia population of black-backed South African aphids (EuphorbiaSA) (d, d’) collected on *Euphorbia caput-medusae*, the natural host species of EcmLV. The lengths of the processus terminalis, the base of the last antennal segment, the siphunculi and the cauda were measured in *A. craccivora* specimens (b, c) and in EuphorbiaSA specimens (e, f). (g) shows the absence of rhinaria on the IVth antennal segment of an alate EuphorbiaSA aphid. Based on these discriminating morphological features, EuphorbiaSA was identified as a member of the formerly described *Aphis tirucallis* (Blackman and Eastop, 2000).

**Supplemental Table 1.** Description of aphid species and populations used in this study and associated references.

**Supplemental Table 2.** Description of aphid species and populations mentioned in this study and associated references.

