## Supplemental Materials for "Alfalfa leaf curl virus is transmitted by *Aphis craccivora* in a highly specific circulative manner"

CIRAD, UMR BGPI, Montpellier, France

BGPI, INRA, CIRAD, Montpellier SupAgro, Univ Montpellier, Montpellier, France

### Supplemental Materials

**Supplemental Figure 1.** Morphological comparisons between specimens of a population of *A. craccivora* sensu stricto (a, a') and specimens of a euphorbia population of black-backed South African aphids (EuphorbiaSA) (d, d') collected on Euphorbia caput-medusae, the natural host species of EcmLV.

**Supplemental Table 1.** Description of aphid species and populations used in this study and associated references.

**Supplemental Table 2.** Description of aphid species and populations mentioned in this study and associated references.

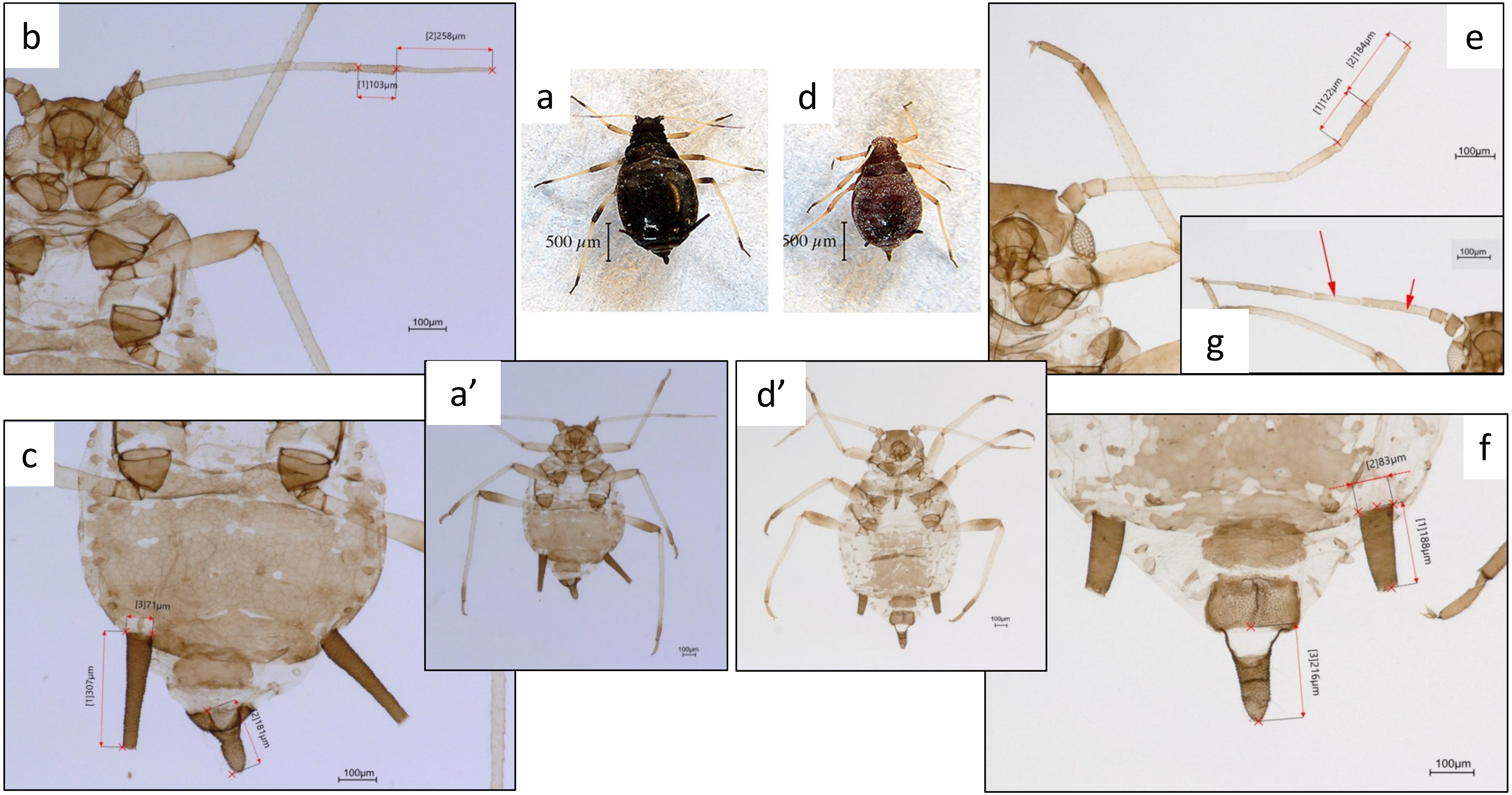


**Supplemental Figure 1.** Morphological comparisons between specimens of a population of *A. craccivora sensu stricto* (a, a’) and specimens of a euphorbia population of black-backed South African aphids (EuphorbiaSA) (d, d’) collected on *Euphorbia caput-medusae*, the natural host species of EcmLV. The lengths of the processus terminalis, the base of the last antennal segment, the siphunculi and the cauda were measured in *A. craccivora* specimens (b, c) and in EuphorbiaSA specimens (e, f). (g) shows the absence of rhinaria on the IVth antennal segment of an alate EuphorbiaSA aphid. Based on these discriminating morphological features, EuphorbiaSA was identified as a member of the formerly described *Aphis tirucallis* (Blackman and Eastop, 2000).

**Supplemental Table 1.** Description of aphid species and populations used in this study and associated references.

| **Species** | **Clone or population** | **Sampling** | | | | | | **Genbank accession** | **BOLD Number** |
|---|---|---|---|---|---|---|---|---|---|
| | | **Country** | **Locality** | **Date** | **Host plant** | **Sampler** | **Reference** | | |
| *Acyrthosiphon pisum* (Harris, 1776) | LL01 | France | Lusignan | 1988 | *Medicago sativa* L.<br>Alfalfa | R. Bournoville (Inra-Lusignan) | Grenier et al. 1994 | | |
| *Aphis craccivora* Koch*, 1854* | EuphorbiaSA | South Africa | Buffelsfontein Game and Nature Reserve | 2015 | *Euphorbia caput-medusae* L. | P. Roumagnac (Cirad-Montpellier) | Roumagnac et al. 2015 | | |
| *Aphis craccivora* Koch*, 1854* | Medicago | France | Montferrier-sur-Lez | 2018 | *Medicago sativa* L.<br>Alfalfa | N. Sauvion & F. Ryckebusch (Inra-Montpellier) | this study | | |
| *Aphis craccivora* Koch*, 1854* | Robinia | France | Prades-le-Lez | 2015 | *Robinia pseudoacacia* L.<br>Common Acacia | G. Labonne (Inra-Montpellier) | this study | | |
| *Aphis craccivora* Koch*, 1854* | Vicia | France | Montferrier-sur-Lez | 2018 | *Vicia sativa* L.<br>Common Vetch | N. Sauvion & F. Ryckebusch (Inra-Montpellier) | this study | | |
| *Aphis fabae* Scopoli, 1763 | A06-405 | Switzerland | St. Margrethen | 2006 | *Chenopodium album* L.<br>Baconweed | C. Vorburger | Vorburger et al. 2017 | | |
| *Aphis fabae* Scopoli, 1763 | AFCOL | France | Colmar | 1972 | *Vicia faba* L. cv. Séville<br>Broad bean | Y. Bouchery (Inra-Colmar) | Kozłowska-Makulska et al. 2009 | | |
| *Aphis gossypii* Glover, 1877 | NM1 | France | Navacelles | 1988 | *Curcubita maxima* Duschene<br>Squash | G. Labonne (Inra-Montpellier) | Lupoli et al.1992<br>Carletto et al. 2009 | | |
| *Dysaphis* (Pomaphis) *plantaginea* (Passerini, 1860) | Nov08 | France | Noves | 2008 | *Malus communis* L. cv. Golden Delicious | M. Siegwaert (Inra-Avignon) | this study | | |
| *Myzus* (Nectarosiphon) *persicae* (Sulzer, 1776) | Lav85 | France | Lavérune | 1985 | *Convolvulus arvensis* L.<br>Bearwind | G. Labonne (Inra-Montpellier) | Labonne et al. 1994 | | |
| *Therioaphis trifolii* (Monell, 1882) | Mon18 | France | Montferrier-sur-Lez | 2018 | *Medicago sativa* L.<br>Alfalfa | M. Perterschmitt, M. Garnier, F. Ryckebusch (Inra-Montpellier) | this study | | |

**Supplemental Table 1** (continuation)

| Species | Group | Clone or population | Experiments | | | Rearing conditions | | | |
|---|---|---|---|---|---|---|---|---|---|
| | | | Transmission tests | Persistence specificity | Molecular analysis | Host plant | Containment | Temperature | Photoperiod |
| *Acyrthosiphon pisum* (Harris, 1776) | | LL01 | ALCV | X | X | *Vicia faba* L. cv. Aquadulce Broad bean | P2 | L24:D21°C | L16:D8h |
| *Aphis craccivora* Koch, *1854* | *A. craccivora* | EuphorbiaSA | ALCV, EcmLV | X | X | *Vicia faba* L. cv. Robinhood Broad bean | P3 | 24± 2°C | L14:D10h |
| *Aphis craccivora* Koch, *1854* | *A. craccivora* | Medicago | ALCV | | X | *Vicia faba* L. cv. Séville Broad bean | P2 | L24:D21°C | L16:D8h |
| *Aphis craccivora* Koch, *1854* | *A. craccivora* | Robinia | ALCV | X | X | *Vicia faba* L. cv. Séville Broad bean | P2 | L24:D21°C | L16:D8h |
| *Aphis craccivora* Koch, *1854* | *A. craccivora* | Vicia | ALCV | | X | *Vicia faba* L. cv. Séville Broad bean | P2 | L24:D21°C | L16:D8h |
| *Aphis fabae* Scopoli, 1763 | | A06-405 | ALCV | | | *Vicia faba* L. cv. Séville Broad bean | P2 | L24:D21°C | L16:D8h |
| *Aphis fabae* Scopoli, 1763 | | AFCOL | ALCV | | X | *Vicia faba* L. cv. Séville Broad bean | P2 | L24:D21°C | L16:D8h |
| *Aphis gossypii* Glover, 1877 | | NM1 | ALCV, PILV, EcmLV | | X | *Cucurbita pepo* L. cv. Diamant Pumpkin | P2 | L24:D21°C | L16:D8h |
| *Dysaphis* (Pomaphis) *plantaginea* (Passerini, 1860) | | Nov08 | EcmLV, PILV | | X | *Plantago lanceolata* L. Buckhorn Plantain | P2 | 23± 4°C | L14:D10h |
| *Myzus* (Nectarosiphon) *persicae* (Sulzer, 1776) | | Lav85 | ALCV, EcmLV | | X | *Solanum melongena* L. Aubergine | P2 | L24:D21°C | L16:D8h |
| *Therioaphis trifolii* (Monell, 1882) | | Mon18 | ALCV | | X | *Medicago sativa* L. Alfalfa | P2 | L26:D24°C | L16:D8h |

**Supplemental Table 2.** Description of aphid species and populations mentioned in this study and associated references.

| Species | Group | Genbank accession | BOLD Number | Sampling | | | | | | Reference |
|---|---|---|---|---|---|---|---|---|---|---|
| | | | | Date | Country | Region | Locality | Host plant | Sampler | |
| *Acyrthosiphon* (Acyrthosiphon) *pisum* (Harris, 1776) | | FJ411411 | | 1999 | USA | Wisconsin | Madison | *Medicago lupulina* L. [Fabaceae] | | Brady et al. 2013 |
| *Aphis* (Aphis) *coronillae* Ferrari, 1872 | *A. craccivora* | | ACEA218-14 | 1999 | France | Alsace | Kaysersberg | | A. Coeur d`Acier | Cœur d'Acier et al. 2014 |
| *Aphis* (Aphis) *coronillae* Ferrari, 1872 | *A. craccivora* | KF638762 | ACOE1245 | 2001 | France | Auvergne | Chouvigny | *Trifolium sp.* L. [Fabaceae] | A. Coeur d`Acier | Cœur d'Acier et al. 2014 |
| *Aphis (Aphis) craccivora* Koch, *1854* | *A. craccivora* | KF362035 | | | Algeria | Ghardaia | | *Medicago sativa* L. [Fabaceae] | | NCBI |
| *Aphis (Aphis) craccivora* Koch, *1854* | *A. craccivora* | | CNBAN433-13 | 2011 | Australia | Australian Capital Territory | | unknown (Malaise traps) | P. Hebert | NCBI |
| *Aphis (Aphis) craccivora* Koch, *1854* | *A. craccivora* | MH571742 | | 2018 | Bangladesh | | | | | Foottit et al. 2008 |
| *Aphis (Aphis) craccivora* Koch, *1854* | *A. craccivora* | EU701313 | | 1995 | Canada | New Brunswick | | | | Song et al. 2016 |
| *Aphis (Aphis) craccivora* Koch, *1854* | *A. craccivora* | KT889380 | | 2015 | China | Henan | Zhengzhou | Locust trees [Fabaceae] | | NCBI |
| *Aphis (Aphis) craccivora* Koch, *1854* | *A. craccivora* | | GMESD081-14 | 2013 | Egypt | Alexandria | Smouha | | O.El-Ansary | Cœur d'Acier et al. 2014 |
| *Aphis (Aphis) craccivora* Koch, *1854* | *A. craccivora* | | ACEA536-14 | 2002 | Greece | Korinthia (el) | Peloponnese | | | Foottit at al. 2008 |
| *Aphis (Aphis) craccivora* Koch, *1854* | *A. craccivora* | EU701309 | RDBA768-06 | 2006 | Hawaii | Maui | Kihei | | R.G. Foottit, R. Miller, K.S. Pike | NCBI |
| *Aphis (Aphis) craccivora* Koch, *1854* | *A. craccivora* | KR017752 | | | India | | | | | Komazaki et al. 2010 |

| Species | Group | Genbank accession | BOLD Number | Sampling | | | | | | Reference |
|---|---|---|---|---|---|---|---|---|---|---|
| | | | | Date | Country | Region | Locality | Host plant | Sampler | |
| *Aphis (Aphis) craccivora* Koch, *1854* | *A. craccivora* | KY323013 | | | Kenya | | | | | Lagos et al. 2014 |
| *Aphis (Aphis) craccivora* Koch, *1854* | *A. craccivora* | KC897559 | GBMHH16383-19 | 2009 | Madagascar | | | *Vigna unguiculata* (L.) Walp. [Fabaceae] | M. Randrianandrasana | Ashfaq et al. 2018 |
| *Aphis (Aphis) craccivora* Koch, *1854* | *A. craccivora* | KY838020 | | 2012 | Pakistan | Islamabad | Shakarparian forest | unknown (Malaise traps) | M. Rafique | NCBI |
| *Aphis (Aphis) craccivora* Koch, *1854* | *A. craccivora* | | CGPTA021-09 | 1989 | Turkey | Erzurum | Horasan | | S. Guclu | Foottit et al. 2008 |
| *Aphis (Aphis) craccivora* Koch, *1854* | *A. craccivora* | GU668252 | | 1999 | USA | Colorado | | | | NCBI |
| *Aphis (Aphis) craccivora* Koch, *1854* | *A. craccivora* | HQ971366 | | 2009 | Argentina | Neuquen | Cutral-Co | *Medicago sativa* L. [Fabaceae] | Perez Hildago, N. | Pilar et al. 2012 |
| *Aphis* (Aphis) *cytisorum* Hartig, 1841 | | HQ971362 | | 2009 | Argentina | Río Negro | Santuario Virgen Misionera | *Cytisus scoparius* L. [Fabaceae] | Perez Hildago, N. | Pilar et al. 2012 |
| *Aphis* (Aphis) *cytisorum* Hartig, 1841 | | | ACEA245-14 | 2000 | France | Occitanie | Anduze | | | BOLD |
| *Aphis* (Aphis) *fabae* mordvilkoi Börner & Janisch, 1922 | | NC_039988 | | 2017 | Belarus | | Minsk | *Philadelphus sp.*[Hydrangeaceae] | | NCBI |
| *Aphis* (Aphis) *gossypii* Glover, 1877 | | NC_024581 | | | China | | | *Gossypium sp.*, [Malvaceae] | | NCBI |
| *Aphis* (Aphis) *intybi* Koch, 1855 | | | ACEA527-14 | 2002 | France | Occitanie | Montferrier-sur-Lez | | A. Cœur d'Acier | BOLD |
| *Aphis* (Aphis) *intybi* Koch, 1855 | | JX438175 | RFQC247-11 | 2010 | Canada | Québec | Montreal | | Pilon, Claude | Pilar et al. 2012 |

| Species | Group | Genbank accession | BOLD Number | Sampling | | | | | | Reference |
|---|---|---|---|---|---|---|---|---|---|---|
| | | | | Date | Country | Region | Locality | Host plant | Sampler | |
| *Aphis* (Aphis) *rumicis* Linnaeus, 1758 | | | ACEA088-14 | 1997 | France | Seine-Maritime | Criquetot-sur-Ouville | *Rumex sp.* L. [Polygonaceae] | Coeur d'Acier | Cœur d'Acier et al. 2014 |
| *Aphis* (Aphis) *rumicis* Linnaeus, 1758 | | | ACEA312-14 | 2000 | France | Brittany | Fouesnant | *Rumex sp.* L. [Polygonaceae] | Coeur d'Acier | Cœur d'Acier et al. 2014 |
| *Aphis* (Aphis) *spiraecola* Patch, 1914 | *A. craccivora* | JX844405 | | | China | | | | | |
| *Aphis (Aphis) tirucallis* Hille Ris *Lambers, 1954* | *A. craccivora* | KF639083 | ACEA181-14 | 1999 | France | Dordogne | St-Vincent-de-Cosse | *Euphorbia sp.* L. *[Euphorbiaceae]* | A. Cœur d'Acier | Cœur d'Acier et al. 2014 |
| *Dysaphis* (Pomaphis) *plantaginea* (Passerini, 1860) | | KF639366 | | 2005 | France | Languedoc-Roussillon | La Bastide-Puylaurent | *Plantago lanceolata* L. [Fabaceae] | A. Cœur d'Acier & E. Jousselin | Cœur d'Acier et al. 2014 |
| *Dysaphis* (Pomaphis) *plantaginea* (Passerini, 1860) | | KF639368 | | 2008 | France | Languedoc-Roussillon | Gaujac | *Malus domestica* Borckh. [Rosaceae] | A. Cœur d'Acier | Cœur d'Acier et al. 2014 |
| *Dysaphis* (Pomaphis) *reaumuri* (Mordvilko, 1928) | | KF639375 | | 2002 | Greece | Lakonia (el) | Lagada | *Rosaceae sp.* L. | A. Cœur d'Acier | Cœur d'Acier et al. 2014 |
| *Myzus* (Nectarosiphon) *persicae* (Sulzer, 1776) | | KU236024 | | | China | | | | | NCBI |
| *Rhopalosiphum maidis* (Fitch, 1856) | | | ACEA794-14 | 2006 | Italia | Sicily | Catania | | | Cœur d'Acier et al. 2014 |
| *Rhopalosiphum padi* (Linnaeus, 1758) | | | ACEA957-14 | 2008 | France | PACA | Antibes | | | Cœur d'Acier et al. 2014 |
| *Schizaphis* (Schizaphis) *graminum* (Rondani, 1852) | | AY531391 | | | | | | | | Thao et al. 2004 |
| *Sitobion* (Sitobion) *avenae* (Fabricius, 1775) | | KJ742384 | | | China | | | | | Zhang et al. 2016 |
